# Immunofluorescence quality of human brain tissue fixed with solutions used in gross anatomy laboratories

**DOI:** 10.64898/2026.03.04.709624

**Authors:** Eve-Marie Frigon, Victoria Perreault, Amy Gérin-Lajoie, Denis Boire, Josefina Maranzano

## Abstract

Brain banks provide small tissue samples fixed in neutral-buffered-formalin (NBF), but human anatomy teaching laboratories could provide full brains fixed with solutions that are more appropriate for gross anatomy such as a saturated salt solution (SSS) or an alcohol-formaldehyde solution (AFS). Advanced aging and prolonged exposure to aldehydes are known to enhance brain tissue autofluorescence (AF), limiting the efficacy of immunofluorescence (IF) procedures. We have previously shown by IF staining the antigenicity preservation in mouse brains fixed with the three solutions. We now aimed to compare the quality of IF staining in human brains fixed with SSS, AFS and NBF. In addition, we compared the efficiency of AF quenching methods, namely the application of SudanBlackB (SBB) and the treatment of sections with sodium borohydride (NaBH_4_).

Blocks of neocortex were extracted from 18 brains (NBF=6, SSS=6, AFS=6) and cut into 40µm sections. Neurons (anti-NeuN, AlexaFluor-488) and astrocytes (anti-GFAP, AlexaFluor-555) were revealed with IF after an antigen retrieval protocol, while two treatments (SBB or NaBH_4_) were used to quench AF. We then assessed the degree of AF (criteria: background or cell AF) and the immunostaining quality with excitation wavelengths of 488nm, 555nm and 647nm.

Brains fixed with all three solutions showed well-labeled astrocytes, whereas neurons weren’t always stained, but this was not associated to the fixative solution. The overall AF intensity was similar in sections from brains fixed with all three solutions. Finally, the SBB treatment was the most effective at reducing AF in all specimens.

Given the similarity in AF and antigenicity assessment across the three solutions, we conclude that brains fixed with SSS and AFS could be good alternatives for NBF-fixed specimens in the context of IF experiments processed with a SBB protocol.

**Highlights:**

- Immunofluorescence staining is feasible in brains fixed with anatomy labs solutions
- GFAP is less affected by fixation than NeuN
- Autofluorescence can be reduced by Sudan Black treatment

## 1. Introduction

Histology is the gold standard method to visualize cellular structure in both the normal and pathological brain [1-3]. Neuroscientific research therefore requires access to cerebral tissue, which can be obtained through brain banks. However, these banks can usually only provide researchers with a limited number of small tissue blocks (i.e., brains dissected in 1cm^3^ blocks, or at most, single hemispheres) [4]. Furthermore, the standard practice of brain banks is to immerse and store brains in neutral-buffered formalin (NBF) for lengthy periods ranging from years to decades [5]. This extended exposure to aldehydes results in increased protein cross-linking and a reduced preservation of antigenicity [6, 7]. Alternatively, gross anatomy laboratories could be a source of whole brains, making research requiring complete hemispheres or even entire brains possible (e.g., studies of magnetic resonance imaging (MRI), brain connectivity, etc.). However, these specimens are preserved using solutions that differ from the NBF commonly used in brain banking, such as a saturated salt solution (SSS) [8] or an alcohol-formaldehyde solution (AFS) [9]. Hence, these solutions could alter the brain cells’ morphology and antigenicity preservation [10, 11].

Few studies have investigated the effects of these different fixatives on the cellular and molecular characteristics of brain tissue. Previous work of our team has shown that the quality of histochemistry (HC), immunohistochemistry (IHC) and antigenicity of the four main brain cell populations, were preserved in brains fixed with the three solutions, although of lower quality in brains fixed with SSS [9]. Other existing research comparing different fixative solutions has mainly focused on MRI studies [12, 13], microbiological analyses [14], and studies using mouse or rat brains [15, 16], or even other organs [17, 18]. Therefore, the suitability of these solutions for the preservation of human brain tissue for different techniques currently applied in neuroscientific research has not been thoroughly assessed, especially for gold-standard histology techniques.

Immunofluorescence (IF) allows for the visualization of a wide variety of antigens through its use of fluorophores [19]. IF offers numerous advantages in its applications, namely signal amplification by indirect labeling [20]. It also allows for multiplex staining to assess antigen colocalization using different fluorophores [19] and can allow for non-destructive 3D imaging spatial relationships between molecular and cellular labels [21]. Since labeled cells are observed against an unmarked black background, confocal microscopy can be used to acquire multiplanar images allowing high-quality tridimensional cellular reconstructions, ideal for studying cell morphology [22]. Even though IF staining is not widely used in neuropathology assessment because of its sensitivity to photobleaching [23], IF remains a method of choice in neuroscientific research [24-26]. A recent study assessed the quality of IF in human brains fixed with NBF to optimize fixation procedures and define acceptable post-mortem intervals [7]. Moreover, we previously validated the use of IF staining for neurons and astrocytes in mouse brains fixed with NBF, SSS and AFS [16]. However, the quality of IF in post-mortem human brain tissue preserved with alternate fixatives has yet to be clearly assessed.

Despite the numerous advantages of IF staining, autofluorescence (AF) artifacts can make this technique particularly challenging. Indeed, nonspecific background autofluorescence and cellular elements such as lipofuscin are present in elderly brains, which are most prevalent in body donor programs [27]. For example, the study mentioned above [7] reported observing lipofuscin-like AF, but they did not assess this artifact in their analyses. AF artifacts are also produced by free aldehydes remaining in the tissue, which react with protein amine groups to form Schiff bases, known to produce AF [28, 29]. The autofluorescence can interfere with the specific signal of IF-labeled antigens and hinder microscopic analysis [30]. Many IF studies address this issue by employing various AF-quenching treatments, such as photobleaching, masking with Sudan Black B (SBB) or sodium borohydride (NaBH_4_) [31, 32]. While photobleaching may not only affect AF but also reduce the specific labeling of cells of interest [23], SBB and NaBH_4_ are methods of choice [31].

The overall goal of this study was therefore to compare the quality of IF labeling performed on brains fixed with SSS and AFS, as standard practice of gross anatomy laboratories, to that of brains fixed with NBF, typically used in brain banking. More specifically, we assessed the quality of IF staining of neurons and astrocytes, as well as AF associated with each fixation method. Furthermore, we tested the efficiency of two widely used AF quenching treatments – SBB and NaBH_4_ – to optimize IF protocols for human brains fixed with solutions commonly used in gross anatomy laboratories.

## 2. Methods

### 2.1 Population

We used a convenience sample of 18 specimens fixed by perfusion with SSS (N=6), AFS (N=6) and NBF (N=6) from the body donation program of the Université du Québec à Trois-Rivières (UQTR). Prior to their death, donors consented to the donation of their body for teaching and research purposes, and the sharing of their anonymized medical information. Finally, this study was approved by UQTR’s Ethic Subcommittee on Anatomical Teaching and Research. The donor’s characteristics are reported in Table 1.

**Table 1:** Demographic variables of the specimens.

| ID | Fixative | Sex | Age | Cause of death | PMI (hours) | FL (years) | Histology block of neocortex |
| --- | --- | --- | --- | --- | --- | --- | --- |
| 1 | NBF | F | 84 | Sepsis, acute pyelonephritis | 21 | 5 | Occipital |
| 2 |  |  | 89 | Pneumonia | 23 | 4 | Frontal |
| 3 |  |  | 44 | Ovarian cancer | 45 | 6 | Temporal |
| 4 |  | M | 61 | Pulmonary neoplasia | 24 | 12 | Frontal |
| 5 |  |  | 85 | Pulmonary edema | 12 | 8 | Frontal |
| 6 |  |  | 74 | Pulmonary neoplasia | 3 | 4 | Frontal |
| Mean |  | 3F - 3M | 72.8±17.4 | N/A | 21.3±14.1 | 6.5±3.1 | N/A |
| 7 | SSS | F | 67 | COPD | 36 | 5 | Temporal |
| 8 |  |  | 84 | Myocardial infarction | 60 | 4 | Occipital |
| 9 |  | M | 87 | Lung cancer | 7,5 | 6 | Parietal |
| 10 |  |  | 83 | Heart failure | 20 | 4 | Parietal |
| 11 |  |  | 81 | Colon cancer | 26 | 3 | Frontal |
| 12 |  |  | 78 | Cholangiocarcinoma | 39 | 2 | Frontal |
| Mean |  | 2F-4M | 80±7.0 | N/A | 31.4±18.0 | 4±1.4 | N/A |
| 13 | AFS | F | 64 | Pneumonia, dementia | 15,5 | 5 | Temporal |
| 14 |  |  | 96 | Breast neoplasia | 13 | 3 | Frontal |
| 15 |  |  | 80 | Peritoneal carcinomatosis | 23 | 3 | Temporal |
| 16 |  |  | 83 | Pancreatic cancer | 18 | 2 | Temporal |
| 17 |  | M | 61 | Pulmonary neoplasia | 24 | 12 | Temporal |
| 18 |  |  | 91 | COPD | 20 | 3 | Parietal |
| Mean |  | 4F-2M | 79.2±14.1 | N/A | 18.9±4.3 | 4,7±3.7 | N/A |

NBF=Neutral-buffered formalin; SSS=Saturated salt solution; AFS=Alcohol-formaldehyde solution; F=Female; M=Male; COPD=Chronic obstructive pulmonary disease; PMI=Post-mortem interval; FL=Fixation length; N/A=Not applicable

### 2.2 Histology protocols

#### 2.2.1 Sectioning

Neocortical tissue blocks (about 1 cm^3^) were dissected from each specimen and rinsed in phosphate-buffered saline (PBS) (0.1 M, 0.9% NaCl, pH 7.3) for five minutes. For cryoprotection, blocks were immersed in a 30% sucrose solution in PBS and stored at 4 °C until they sank, indicating complete diffusion of sucrose into the tissue. Blocks were then frozen at −80 °C before cutting at 40 µm using a cryostat (Leica CM1950) at −19 °C. Sections were stored in well plates filled with PBS until further use.

#### 2.2.2 Antigen retrieval

A heat-induced antigen retrieval (HIAR) protocol was applied to all sections to restore the conformation of proteins potentially altered by chemical fixation. Free-floating sections were incubated in boiling 0.01 M citrate buffer (pH 6) for 20 minutes, then immediately transferred to a cold bath of deionized water for 10 minutes to induce thermal shock, facilitating protein conformational changes and reducing folding of the sections [33]. Sections were then stored in PBS until further processing.

#### 2.2.3 Immunofluorescence

Free-floating sections were rinsed in PBS for 5 minutes, then incubated in 20% methanol for 20 minutes to permeabilize cellular membranes. This was followed by three 5-minutes washes (3×5min) in PBS. Sections were then incubated for 2 hours on a shaker in a blocking solution (4% normal donkey serum, 1% bovine serum albumin (BSA), 1% glycerol, and 0.3% Triton X-100 in 0.1M PBS), to prevent nonspecific antibody binding. Sections were then directly transferred into the same blocking solution containing the primary antibodies (Table 2) and incubated overnight at room temperature on a shaker.

**Table 2:** Antibodies used for immunofluorescence staining.

| **Primary antibodies** | **Animal** | **Lab** | **Dilution** | **Code** |
| --- | --- | --- | --- | --- |
| Anti-Neuronal nuclei (NeuN) | Mouse | Abcam | 1:1000 | Ab104224 |
| Anti-Glial fibrillary acidic protein (GFAP) | Rabbit |  |  | Ab68428 |
| **Secondary antibodies** | **Animal** | **Lab** | **Dilution** | **Code** |
| Anti-Mouse Alexa Fluor 488 | Donkey | Abcam | 1:500 | Ab150105 |
| Anti-Rabbit Alexa Fluor 555 |  | NovusBio |  | NBP1-75274 |

Sections were rinsed (3×5min) in PBS, then incubated at room temperature in the dark for 2 hours, in the blocking solution containing the secondary antibodies (Table 2). Following incubation, sections were washed (3×5min) in PBS, then mounted onto gelatin-coated microscope slides (2% gelatin in distilled water) using 0.01 M phosphate buffer (no NaCl). Slides were air-dried in the dark and coverslipped with PVA-DABCO aqueous mounting medium (Millipore Sigma #10981).

Negative controls, in which primary antibodies were omitted during the IF protocol, were processed in parallel of IF-processed sections to verify the specificity of the secondary antibodies used. These controls were also used to assess AF intensity in three different channels (488nm, 555nm, and 647nm).

#### 2.2.4 Treatment

All specimens – including both experimental slides and negative controls – were treated with NaBH_4_ or Sudan Black (separately) to quench AF. The effectiveness of these treatments was evaluated by comparison with untreated counterparts.

##### 2.2.4.1 NaBH_4_

Prior to HIAR and IF protocols, free-floating sections were incubated four times for 15 minutes in a freshly prepared solution of NaBH_4_ (1 mg/mL). A new solution was prepared before each incubation to preserve NaBH_4_ reactivity. Sections were then rinsed three times for 5 minutes in 0.1 M PBS.

##### 2.2.4.2 Sudan Black B

SBB (S-2380, Sigma) solution (0.3% in 70% ethanol) was freshly prepared, stirred with a magnetic stir bar for 15 minutes, and filtered twice through an 11 µm filter. Following the HIAR and IF protocols, mounted and air-dried sections were immersed in the solution for 12 minutes [25]. Slides were then rapidly rinsed eight times with 0.1M PBS and coverslipped using PVA-DABCO aqueous mounting medium (Millipore Sigma #10981).

### 2.3 Microscopy and variables of interest

All microscopic analyses were conducted using a Spinning disk confocal microscope (Olympus BX51W1) controlled by Neurolucida (MBF Bioscience) software. We assessed the following variables by looking at all the sections that were available for each of the specimens and treatments. For each of the three wavelengths, we adjusted the camera parameters (i.e., exposure and sensitivity) consistently. However, the parameters were modified for the sections that were processed with SBB treatment, since the background was a lot darker (the parameters were increased to discern the signal).

#### 2.3.1 Immunofluorescence quality

The homogeneity of cell distribution and completeness of antibody penetration in the sections were the criteria used to assess the quality of IF labeling of neurons and astrocytes in the tissue samples.

##### 2.3.1.1 Antigenicity distribution

We used a low power objective (UPlanSApo 4x/0.16∞/-/FN26.5) to assess the cell distribution over a large field of view and to scan the sections. We scored the specimens as follows:

0 = Absence of labeling (Figure 1A); 1= Isolated cells (Figure 1B); 2 = Cell patches (Figure 1C); 3 = Homogeneous distribution (Figure 1D).

**FIGURE 1.**
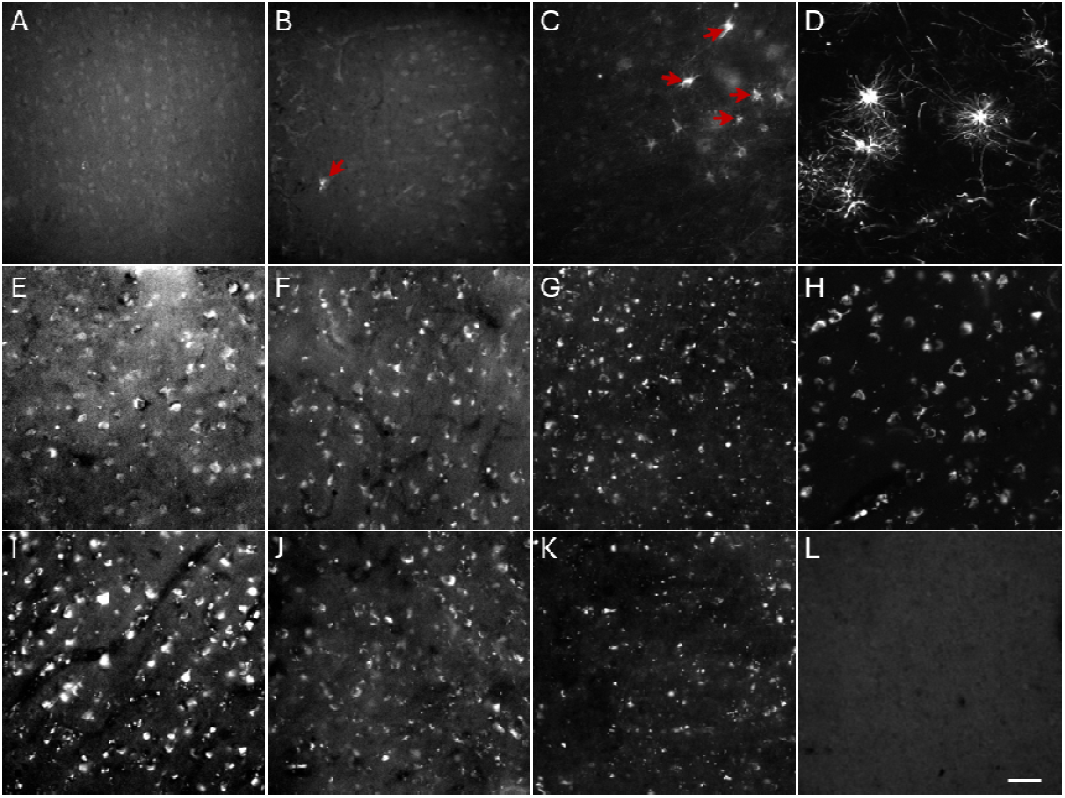
Photomicrographs (20X) illustrating the qualitative scales used to assess antigenicity preservation and AF intensity. Immunofluorescence staining for astrocytes (GFAP) (A-D) showing absence of labeled cells (A), isolated cells (B), cells in patches (C) and homogeneous distribution of cells (D). Unlabeled brain sections showing AF intensity of the background (E-H): bright gray (E), gray (F), dark gray (G) and black (H) backgrounds. Unlabeled brain sections showing AF intensity of the cells (I-L): very high (I), high (J), low (K) and very low (L) cell AF. Scale bar = 50μm.

##### 2.3.1.2 Antibody penetration

The labeling quality is dependent on the degree of penetration of the antibodies into the sections. In free-floating sections, while the antibodies can penetrate through both sides of the section, there can also be an incomplete penetration. This would result in absent or weaker staining in the middle of the section depth, producing an alternate (“sandwich”) labelling pattern. Antibody penetration was scored as follows:

0 = Incomplete; 1 = Complete.

#### 2.3.2 Autofluorescence intensity

We assessed the AF intensity through a low power objective (4X) and a higher magnification (20X, UPlanSApo 20X/0.75∞0.17/FN26.5). We assessed the background AF as it interferes with the cell/background contrast, reducing signal-to-noise ratio. We also assessed the AF of cells, since this could lead to falsely identified immunolabeled cells. Since the AF may be different when the tissue is excited with different wavelengths [34], we assessed AF variables at 488, 555 and 647 nm (in grayscale display). A high intensity of AF is not ideal, as it would obscure labeled cells and hinder the analysis, whereas an absence of AF is the best-case scenario, as the labeled cells can be positively and correctly identified.

We scored background autofluorescence as follows:

0 = Bright gray (Figure 1E); 1 = Gray (Figure 1F); 2 = Dark gray (Figure 1G); 3 = Black (Figure 1H).

We also scored the autofluorescence of cells as follows:

0 = Very high (Figure 1I); 1 = High (Figure 1J); 2 = Low (Figure 1K); 3 = Very low (Figure 1L).

### 2.4 Statistical analyses

We scored the variables presented in section 2.3 to create qualitative categories that were compared for two different factors: the fixative solution (NBF, SSS or AFS) as well as the applied treatment (Untreated, NaBH_4_ and SBB) using Chi-square tests. All statistical analyses were adjusted for multiple comparisons using Bonferroni correction. All statistical analyses were conducted using SPSS statistics software (version 29.0.2.0).

## 3. Results

### 3.1 Immunofluorescence quality

#### 3.1.1 According to the fixatives

We found that GFAP immunolabeling of astrocytes was always preserved and produced a homogeneous distribution of labeled cells (score=3). Also, antibody penetration was always complete in all cases, despite the fixative or treatment. Therefore, we found no significant difference between brains fixed with the three solutions.

Similarly, after a Bonferroni correction, the distribution of NeuN-labeled neurons was not significantly different in brain sections fixed with the three solutions and differently treated for AF (Figure 2A), nor when individually assessing untreated (Figure 2B), NaBH_4_-treated (Figure 2C) and SBB-treated (Figure 2D) sections.

**FIGURE 2.**
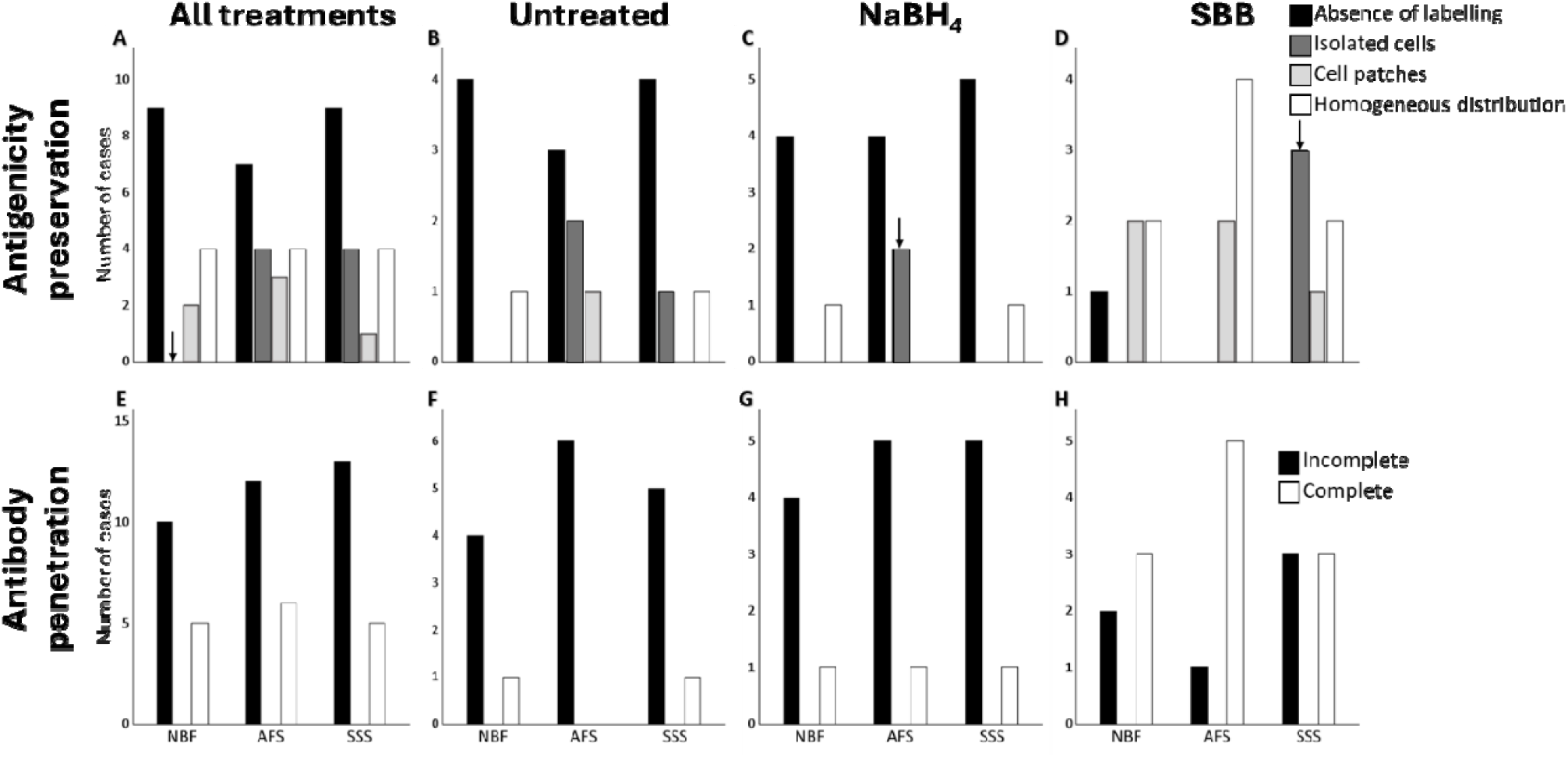
Bar charts showing the immunofluorescence staining quality of neurons (NeuN) in human brain sections treated for autofluorescence according to the fixative solution. Antigenicit preservation in all treated and untreated sections (A), in sections not treated for AF (B), in section treated with NaBH_4_ (C) and in sections treated with SBB (D). Antibody penetration in all treated and untreated sections (E), in sections not treated for AF (F), in sections treated with NaBH_4_ (G) and in sections treated with SBB (H). ↓ = p<0.05, but this did not retain significance after a Bonferroni correction. NBF=Neutral-buffered formalin; AFS=Alcohol-formaldehyde solution; SSS=Saturated salt solution; SBB=Sudan Black B; NaBH_4_=Sodium borohydride.

We also found that the antibody penetration of anti-NeuN was not statistically different in brains fixed with the three solutions, when sections were pooled for all treatments (Figure 2E). In addition, no significant difference was detected in the degree of antibody penetration in brains fixed with the three solutions considering the sections untreated (Figure 2F) or treated with NaBH_4_ (Figure 2G) and SBB (Figure 2H) separately.

#### 3.1.2 According to the treatments

GFAP staining was always homogeneous, and antibody penetration was always complete. Consequently, there was no significant difference in brain sections following the different AF quenching treatments.

The SBB treatment resulted in a significantly lower occurrence of absence of NeuN-labeled neurons (p<0.0001, considering all the fixatives together). Additionally, a higher frequency of homogeneous and patchy distribution of labeled neurons was observed in the sections treated with SBB, but this significant difference did not withstand the Bonferroni correction (Figure 3A). This is because, when fixations procedures are taken separately, we found no SBB-treated sections with an absence of labeling with SSS fixation (significance: p=0.003) (Figure 3D). This was also found in brains with AFS, but it did not retain significance after a Bonferroni correction. However, we found a significantly higher number of sections with a homogeneous distribution following AFS fixation (p=0.001) (Figure 3C).

**FIGURE 3.**
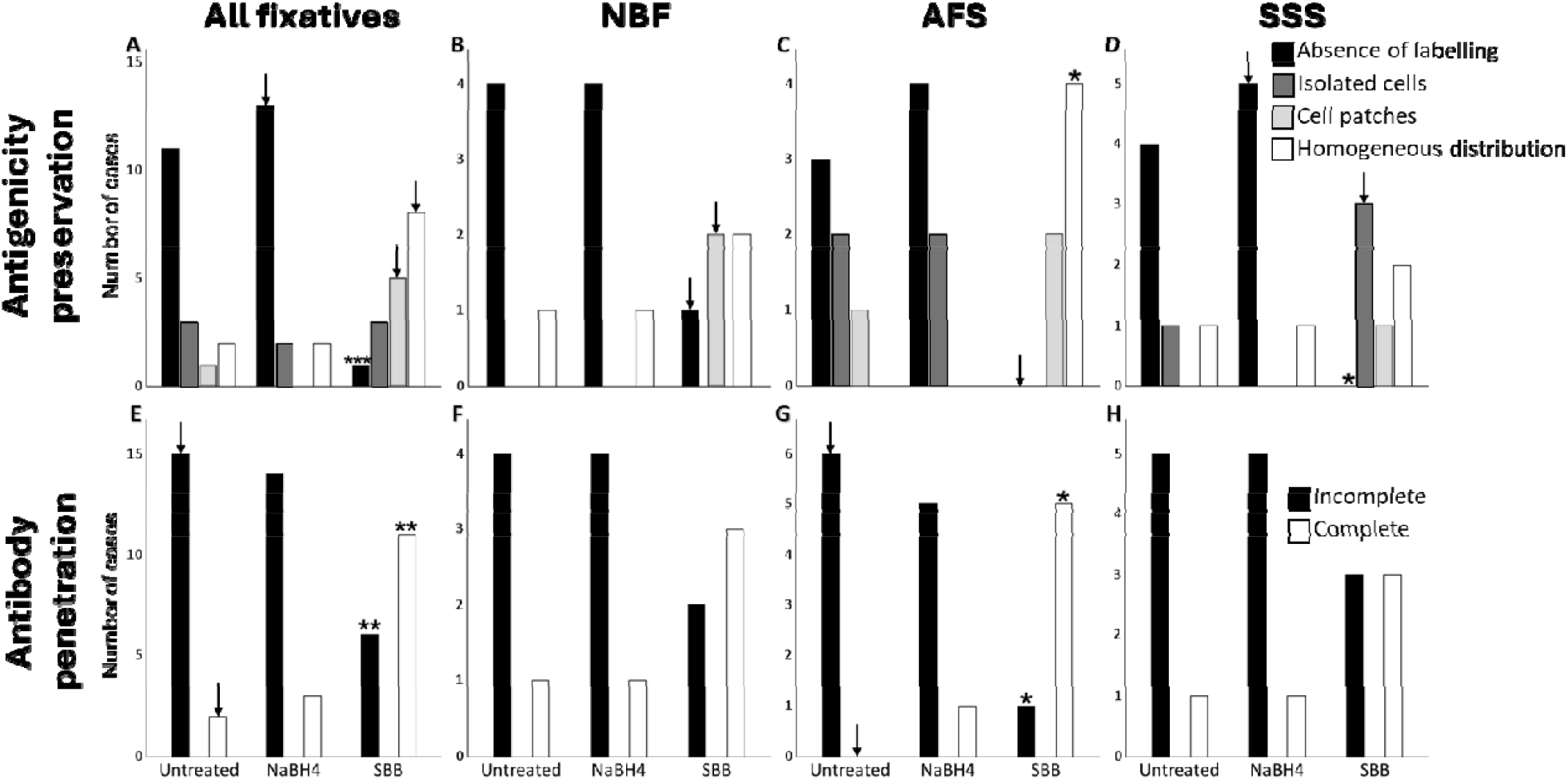
Bar charts showing the immunofluorescence staining quality of neurons (NeuN) in human brain sections fixed with different solutions according to the autofluorescence treatment. Antigenicity preservation in brain sections fixed with all three fixatives (A), in sections fixed with NBF (B), in sections fixed with AFS (C) and in sections fixed with SSS (D). Antibody penetration in brain sections fixed with all three fixatives (E), in sections fixed with NBF (F), in sections fixed with AFS (G) and in sections fixed with SSS (H). ↓ = p<0.05, but this did not retain significance after a Bonferroni correction. For A-D: *p<0.0042; **p<0.001; ***p<0.0001 after a Bonferroni correction. For E-H: *p<0.0083; **p<0.001; ***p<0.0001 after a Bonferroni correction. NBF=Neutral-buffered formalin; AFS=Alcohol-formaldehyde solution; SSS=Saturated salt solution; SBB=Sudan Black B; NaBH_4_=Sodium borohydride.

Furthermore, when considering all fixatives together, we found a significantly higher occurrence of complete antibody penetration and a significantly lower incomplete antibody penetration in SBB-treated sections (p=0.0003) (Figure 3E). This is related to the significantly fewer sections with incomplete penetration and more sections with a complete penetration in brains fixed with AFS (p=0.001) (Figure 3G).

### 3.2 Autofluorescence intensity

#### 3.2.1 Autofluorescence of the background

##### 3.2.1.1 According to the fixative

Background intensity was assessed at different wavelengths as a measure of the level of autofluorescence in sections processed with different autofluorescence mitigation treatments (NaBH_4_ or SBB). In sections of brains fixed with the three fixatives, the highest background intensity was observed with excitation at 488nm. Indeed, brains fixed with all three solutions showed more bright gray backgrounds and fewer black backgrounds at 488nm (Figure 4A) than at 555nm (Figure 4E) or 647nm (Figure 4I) regardless of the applied mitigation treatment. However, we found no significant difference in the background intensity in brains fixed with the three solutions when they were untreated or treated with NaBH_4_ in the different wavelengths. With excitation set to 488nm, we found that the SBB treatment resulted in a higher number of black sections and a fewer number of bright gray sections in brains fixed with NBF (Figure 4D), but this did not retain significance after a Bonferroni correction.

**FIGURE 4.**
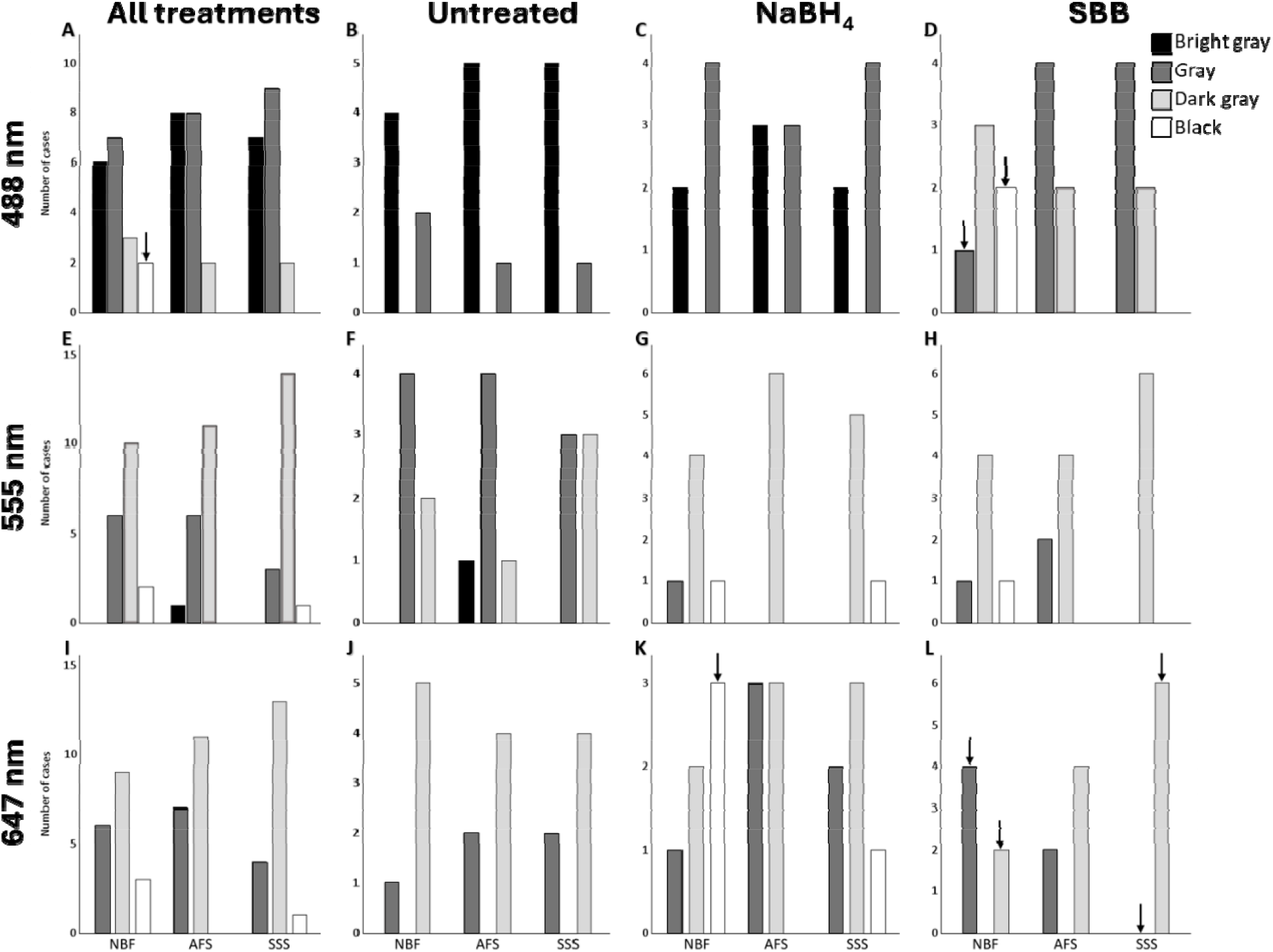
Bar charts showing the background autofluorescence intensity in human brain sections according to the three fixatives. Autofluorescence intensity was measured at 488nm (first row), 555nm (second row) and 647nm (third row) in treated and untreated sections (A, E and I), in untreated sections only (B, F and J), in NaBH_4_-treated sections (C, D and K) and in SBB-treated sections (D, H and L). ↓ = p<0.05, but this did not retain significance after a Bonferroni correction. NBF=Neutral-buffered formalin; AFS=Alcohol-formaldehyde solution; SSS=Saturated salt solution; SBB=Sudan Black B; NaBH_4_=Sodium borohydride.

##### 3.2.1.2 According to the treatments

When considering all fixatives together, with the 488nm excitation wavelength, we found significantly more frequent bright gray backgrounds in untreated sections and significantly less (none) bright gray backgrounds in sections treated with SBB (p<0.0001) (Figure 5A). Furthermore, in sections treated with SBB, we found a higher number of black backgrounds (significant before a Bonferroni correction) and a significantly higher number of dark gray backgrounds (p<0.0001) compared to the sections treated with sodium borohydride or untreated sections (Figure 5A). With the 555nm wavelength, we found a significantly higher number of sections with a gray background (p<0.0001) and fewer number of sections with a dark gray background (p=0.0006) in untreated compared to sections treated with NaBH_4_ or SBB procedures (Figure 5E). Finally, during an excitation with the 647nm wavelength, the NaBH_4_ treatment produced a significantly higher number of sections with black backgrounds (p=0.004) (Figure 5I). This was reflected by the NBF-fixed specimens that also showed more black backgrounds following NaBH_4_ treatment, but this did not retain significance after a Bonferroni correction (Figure 5J).

**FIGURE 5.**
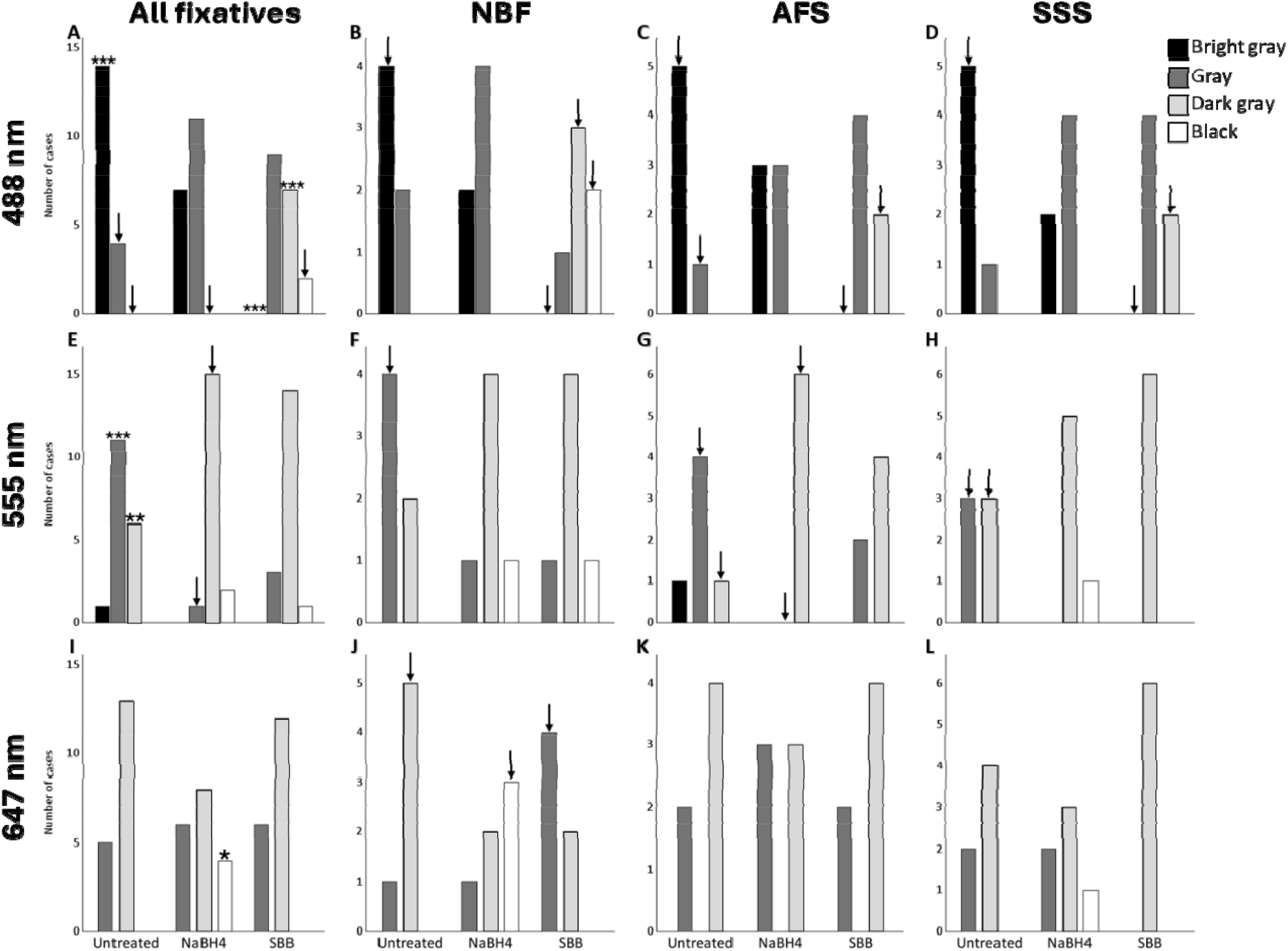
Bar charts showing the background autofluorescence intensity in human brain sections according to autofluorescence quenching treatment. Autofluorescence intensity was measured at 488nm (first row), 555nm (second row) and 647nm (third row) in brain sections fixed with all three solutions combined (A, E and I) and in sections fixed either with NBF (B, F and J), AFS (C, G and K) or SSS (D, H and L). ↓ = p<0.05, but this did not retain significance after a Bonferroni correction. *p<0.0042; **p<0.001; ***p<0.0001 after a Bonferroni correction. NBF=Neutral-buffered formalin; AFS=Alcohol-formaldehyde solution; SSS=Saturated salt solution; SBB=Sudan Black B; NaBH_4_=Sodium borohydride.

#### 3.2.2 Autofluorescence of the cells

##### 3.3.2.1 According to the fixatives

We evaluated the intensity of cell autofluorescence at different wavelengths as a function of the fixation solution for sections treated for AF (NaBH_4_ or SBB) in negative control brain sections, in which the primary antibodies were omitted from the immunolabeling procedure. Cellular AF intensity was higher in the 488nm (Figure 6A) and 555nm (Figure 6E) than at 647nm wavelength (Figure 6I), where the latter showed more very low and low AF intensity in brains fixed with the three solutions. However, we found no significant difference in the cell autofluorescence intensity in brains fixed with the three solutions when they were untreated or treated with NaBH_4_ or SBB in the different wavelengths.

**FIGURE 6.**
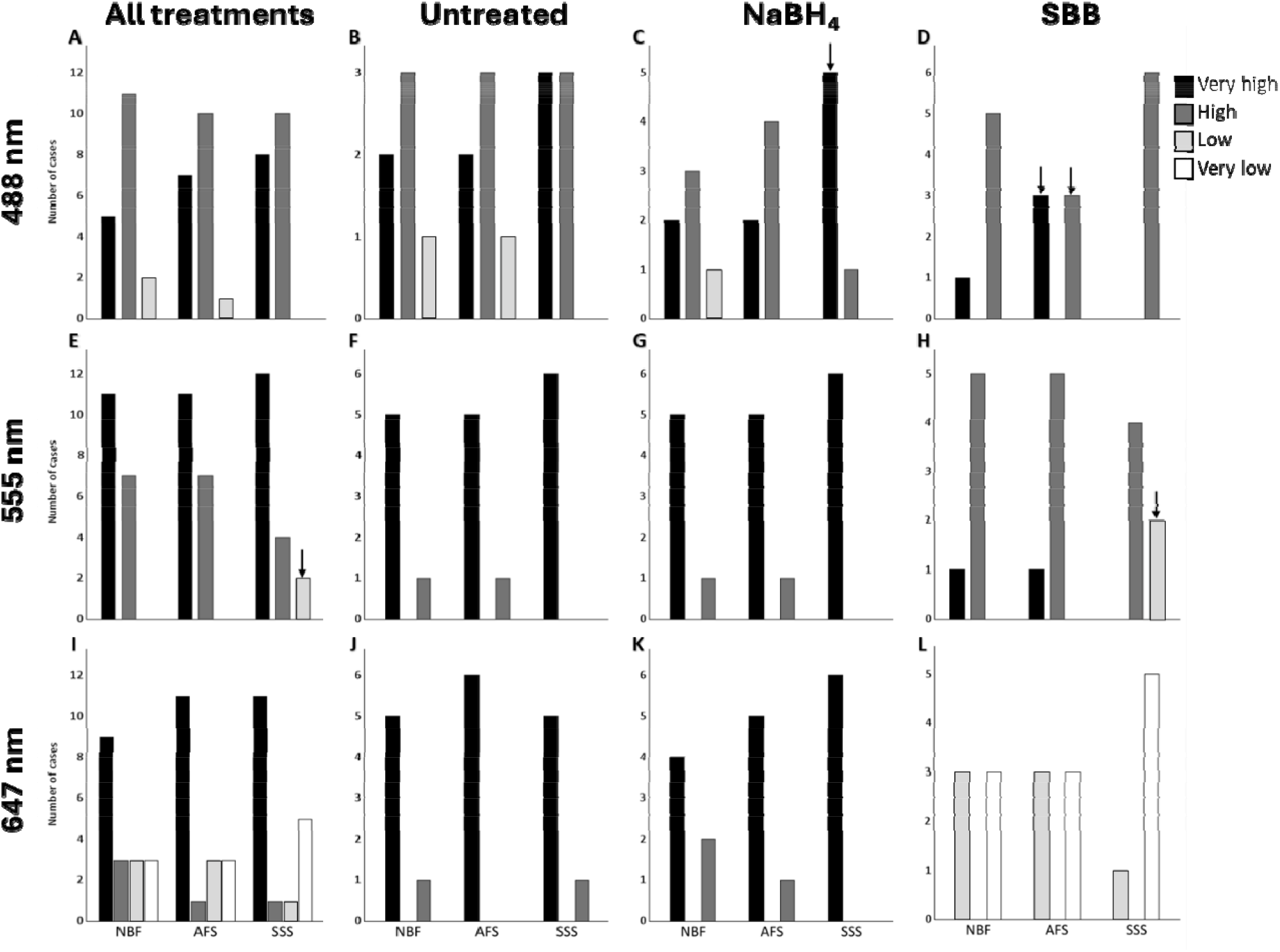
Bar charts showing the autofluorescence intensity of cells in human brain sections according to the three fixatives. Autofluorescence intensity was measured at 488nm (first row), 555nm (second row) and 647nm (third row) in treated and untreated sections (A, E and I), in untreated section only (B, F and J), in NaBH_4_-treated sections (C, G and K) and in SBB-treated sections (D, H and L). ↓ = p<0.05, but this did not retain significance after a Bonferroni correction. NBF=Neutral-buffered formalin; AFS=Alcohol-formaldehyde solution; SSS=Saturated salt solution; SBB=Sudan Black B; NaBH_4_=Sodium borohydride.

#### 3.1.2 According to the treatments

During excitation with the 488 nm wavelength, we found no significant difference in the autofluorescence intensity of cells in untreated or treated (NaBH_4_ or SBB) sections when considering the three fixative groups together (Figure 7A) nor when considered separately (NBF: Figure 7B; SSS: Figure 7C; AFS: Figure 7D). However, we found that there were more sections with high intensity but fewer with very high intensity when processed with a SBB treatment in all fixatives considered together (Figure 7A), or in the SSS-fixed group (Figure 7D), but this did not retain significance after a Bonferroni correction.

**FIGURE 7.**
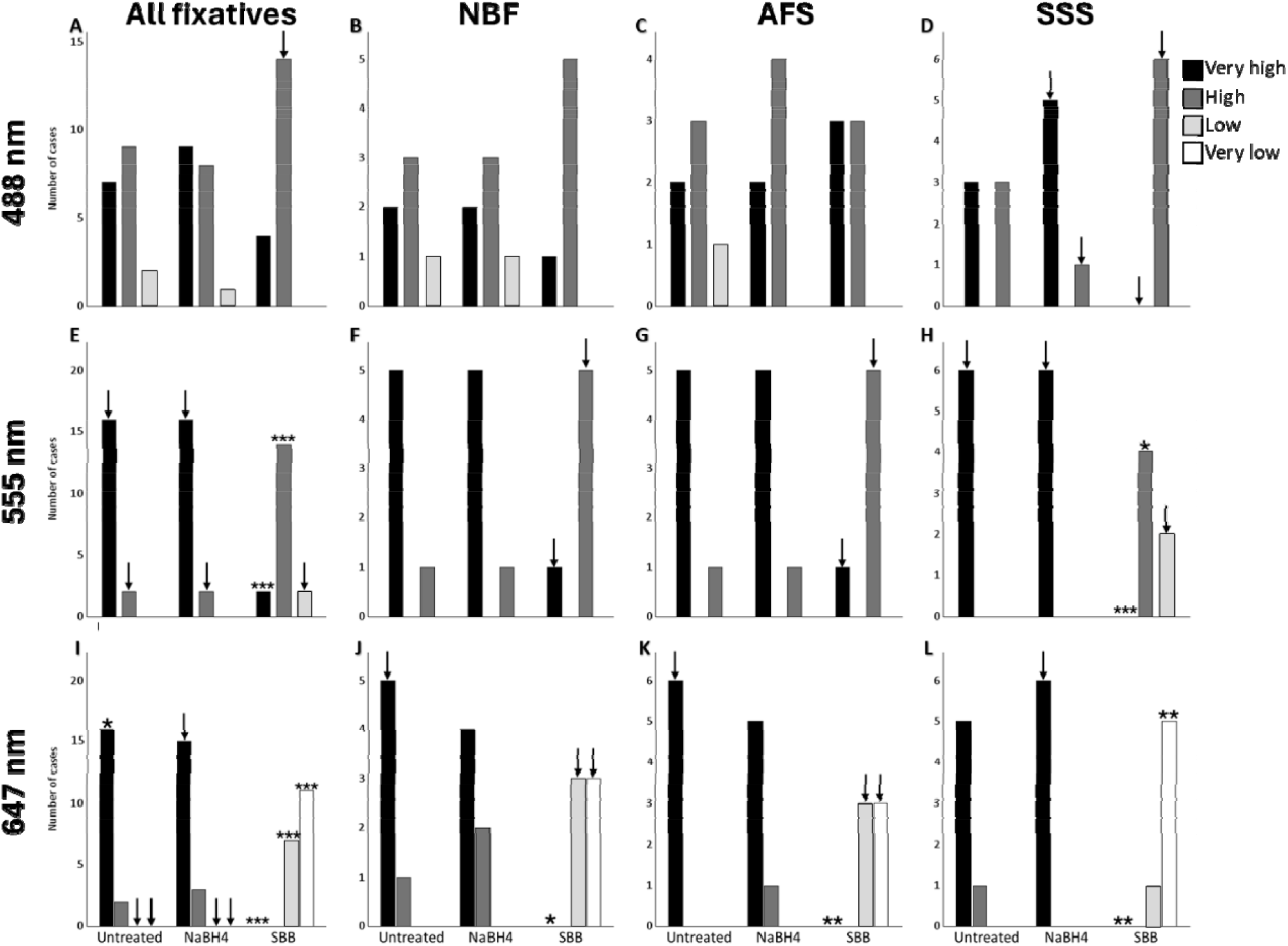
Bar charts showing the autofluorescence intensity of cells in human brain sections according to autofluorescence quenching treatment. Autofluorescence intensity was measured at 488nm (first row), 555nm (second row) and 647nm (third row) in sections fixed with all three solutions combined (A, E and I) and in sections fixed either with NBF (B, F and J), AFS (C, G and K) or SSS (D, H and L). ↓=p<0.05, but this did not retain significance after a Bonferroni correction. *p<0.0042; **p<0.001; ***p<0.0001 after a Bonferroni correction. NBF=Neutral-buffered formalin; AFS=Alcohol-formaldehyde solution; SSS=Saturated salt solution; SBB=Sudan Black B; NaBH_4_=Sodium borohydride.

During excitation with the 555 nm wavelength, when considering all fixatives together, we found a significantly higher number of sections with high AF (p<0.0001) and a lower number of sections with very high AF (p<0.0001) in the SBB treatment group than in the untreated or NaBH_4_-treated group (Figure 7E). The latter showed more sections with very high cellular AF intensity, but this did not retain significance after a Bonferroni correction. Indeed, more sections with high intensity and fewer with very high intensity were found in the SBB group in the NBF-(Figure 7F) and AFS-(7G) fixed specimens, even if these did not retain significance after a Bonferroni correction. Furthermore, when considering the SSS-fixed group, we found a significantly lower number of sections with very high cellular AF (p<0.0001) and a significantly higher number of sections with high cellular AF (p=0.001) in sections treated with SBB (Figure 7H).

Finally, in the 647 nm wavelength, we found a higher number of sections with very high cell AF when they were untreated (p=0.001), while significantly fewer sections with very high cell AF and more sections with low and very low AF intensity of the cells (p<0.0001) were found in the SBB treatment group (Figure 7I). This was reflected by the absence of very high AF in the SBB treatment group for sections fixed with any of the three solutions (NBF: p=0.003 (Figure 7J); AFS: p=0.0001 (Figure 7K) and SSS: p=0.0001 (Figure 7L)). Finally, the SBB treatment group showed a significantly higher number of sections with very low cellular autofluorescence compared to the other two groups in the brains fixed with SSS (p=0.0002) (Figure 7L).

## 4. Discussion

### 4.1 Immunofluorescence quality

Our goal was to assess the quality of immunofluorescent labeling of neurons and astrocytes in brains fixed with NBF from brain banks and two solutions (i.e., SSS and AFS) from anatomy laboratories. These fixatives comprise diverse chemicals that can act distinctly on varied epitopes and alter immunoreactivity [35]. Previous studies have shown that IF labeling of neurons and astrocytes were preserved in mouse brains fixed with solutions in use in gross anatomy laboratories [16] or in NBF-perfused human brains [7]. Similarly, IHC labeling of neurons, astrocytes, microglia and oligodendrocytes was shown in human brains fixed with solutions used in gross anatomy laboratories [9]. We therefore expected that the antigenicity preservation of neurons and astrocytes would be sufficient following an IF protocol in brains fixed with all three solutions.

Our results show that GFAP labeling of astrocytes was always completely preserved, producing a homogeneous distribution of labeled cells with an excitation wavelength of 555 nm in specimens fixed with the three solutions, regardless of the AF mitigation treatment that was applied. This is commensurate with previous IHC studies that showed that, even when confounding variables are inevitable, such as the PMI [7] or even prolonged post-fixation [6], GFAP is strongly immunolabeled.

We also wanted to assess the extent of antibody penetration since incomplete antibody penetration in the sections would preclude 3D cellular reconstructions, which are of interest in cell morphology analysis done with IF [36-38]. We found that anti-GFAP penetrated completely in all specimens fixed with different solutions and with different AF treatments. These results open the door to the use of brains from anatomy laboratories for studies involving astrocytes, such as analyses that differentiate activated vs non-activated astrocytes based on cell morphology [39, 40], or for studying their configuration in synapses [41-43].

However, NeuN IF labeling was completely absent and antibody penetration was incomplete in most of the brain sections that were fixed with the three solutions, reflecting no differences between the chemical fixatives. These results are very different from observations made in mouse brains fixed the three solutions, as every specimen processed showed homogeneous NeuN labeling [16]. NeuN labeling in fixed post-mortem human brains may have been compromised by confounding variables present that are not applicable to mouse brains. For instance, brains obtained from anatomy laboratories are typically from older donors and therefore contain high levels of lipofuscin accumulated within neuronal cell bodies, a pigment known to cause autofluorescence and interfere with the detection of specific cellular labeling [44]. In addition, these brains are subject to post-mortem intervals (PMI; the delay between death and brain fixation), which can adversely affect tissue quality [45] and immunofluorescence staining [7], whereas mouse brains are generally young and fixed immediately at the time of death. However, human brain sections processed with a SBB treatment showed improved preservation of NeuN antigenicity and antibody penetration compared with untreated and NaBH□-treated sections. Because background AF was most intense at 488 nm compared with 555 nm and 647 nm, we surmise that SBB treatment significantly reduced the AF signal, thereby enabling improved visualization of individual NeuN-labeled cells.

### 4.2 Autofluorescence intensity

We also wanted to assess the autofluorescence intensity, as NBF is known to induce intrinsic fluorescence due to its high formaldehyde concentration (4%) [28]. Furthermore, biological tissues are rich in lipofuscin-like substances, also increasing autofluorescence [37, 44, 46]. We therefore expected that, regarding their lower formaldehyde concentration, AFS (1.4%) and SSS (0.8%) would result in reduced AF when compared to brains fixed with NBF. However, we found no significant difference in the autofluorescence intensity of the cells nor of the background in brains fixed with the three solutions, since autofluorescence was high in all brain sections that were assessed. Therefore, it is crucial to develop protocols for quenching autofluorescence in all brains.

We found that SBB treatment was the most effective at reducing AF, as it is a black dye that coats tissue sections masking both background and non-specific cellular labeling without interfering with the fluorescence emitted by the fluorophores [47]. This treatment has already been shown to be the most effective when applied to human brains fixed with NBF [29, 37]. Our results are also consistent with previous findings that SBB treatment reduced 71.88% of AF in the 488nm, 73.68% of AF in the 555nm, and 76.05% in the 647m wavelengths [31]. NaBH_4_ treatment was expected to help reduce background tissue AF, as it is a chemical agent that binds free aldehydes. It has been widely used to quench AF, especially in tissue samples that were fixed with glutaraldehyde [48, 49]. Since glutaraldehyde is composed of two aldehyde groups, whereas formaldehyde only contains one, the effect of NaNH_4_ might have been less powerful with our NBF-fixed specimens [50]. However, it is unlikely to reduce cellular autofluorescence primarily caused by lipofuscin [44, 51, 52], which was in line with our current findings. Indeed, NaBH_4_ treatment only reduced background autofluorescence in the 647nm wavelength in the brains fixed with NBF.

We therefore suggest using antibodies coupled with fluorophores excitable at high wavelengths (>555nm) for IF staining since this bandwidth seems less polluted with AF. Finally, SBB treatment should always be performed when IF staining is of interest in post-mortem fixed human brains.

### 4.3 Limitations

First, our convenience sample resulted in few and heterogeneous (although all from neocortex) tissue samples, which limited the statistical power of our analyses. The small sample size also precluded the inclusion of postmortem interval or fixation length as covariates as well as the comparison of immersed-(brain banks) and perfused-(anatomy laboratories) fixed brains, in which AF could have differed. Therefore, we conducted a descriptive and qualitative analysis to report protocols that were adapted and developed, and to reduce the possible biases of a small sample size in quantitative assessment. Second, we proceeded to a visual and qualitative assessment of the variables of interest, leaving out quantification of the fluorescence signal, since the staining intensity is proportional to gain and exposure settings of the microscope, which could have been biased by the rater. Finally, our assessment focused on two widely used AF mitigation treatments, while other treatments or a combination of the two methods could have improved the results. While we could have optimized the protocols to further quench AF and improve the IF quality for specimens fixed with each specific solution, our goal was to compare brains fixed with the three different solutions by using the same protocols. Future studies increasing the sample size will inform about optimal AF quenching procedures with each fixative solution.

### 4.4 Conclusion

Our study described immunofluorescence protocols and autofluorescence quenching treatments that would be usable in histopathology assessments, multiplex staining or 3D morphologic cell reconstruction studies. Giving the similarity of the immunofluorescence staining quality of neurons and astrocytes in brains fixed with the three solutions, brains from anatomy laboratories would be suitable for neuroscientific research that uses histology techniques and immunofluorescence protocols. Furthermore, we found that Sudan black B treatment is a powerful method to quench autofluorescence that is inevitable in human aging brains. This is promising for neuroscientists interested in using a higher number of brains from anatomy laboratories.

## Acknowledgements

We would like to thank the incredibly generous body donors and their families for making our project possible. We also acknowledge funding resources and the anatomy laboratory staff of the UQTR for their support and help at the lab.

